# A feedback loop between miRNA-155, Programmed cell death 4 and Activation Protein-1 modulates the expression of miR-155 and tumorigenesis in tongue cancer

**DOI:** 10.1101/394130

**Authors:** Shabir Zargar, Vivek Tomar, Vidyarani Shyamsundar, Ramshankar Vijayalakshmi, Kumaravel Somasundaram, Devarajan Karunagaran

**Author notes:** **Correspondence:** Dr D Karunagaran, Department of Biotechnology, Bhupat and Jyoti Mehta School of Biosciences, Indian Institute of Technology Madras, Adyar, Chennai 600036, India.

## Abstract

miR-155 is an oncomir, generated as a non-coding RNA from *BIC* gene whose promoter activity is mainly controlled via AP-1 and NF-κB transcription factors. We found that the expression levels of miR-155 and Pdcd4 exhibit inverse relationship in tongue cancer cells (SAS and AWL) and tumor tissues compared to normal FBM cells and normal tongue tissues, respectively. Insilco and In-vitro studies with 3’UTR of Pdcd4 via luciferase reporter assays, qPCR and western blots show that miR-155 directly targets Pdcd4 mRNA and blocks its expression. Ectopic expression of Pdcd4 or knockdown of miR-155 in tongue cancer cells predominantly reduces AP-1 dependent transcriptional activity of *BIC* promoter and decreases miR-155 expression. In this study, we demonstrate that miR-155 expression is modulated by a feedback loop between Pdcd4, AP-1 and miR-155 which results in enhanced expression of miR-155 with a consequent progression of tongue tumorigenesis. Further, miR-155 knockdown increases apoptosis, arrests cell cycle, regresses tumor size in xenograft nude mice and reduces cell viability and colony formation in soft agar and clonogenic assays. Thus, the restoration of Pdcd4 levels by the use of molecular manipulation such as using miR-155 sponge have important role in the therapeutic intervention of cancers, including tongue cancer.

## 1. INTRODUCTION

MicroRNAs (miRs) are small non-coding RNAs of 18-25 nucleotides in length involved in post-transcriptional gene regulation mostly by binding to the 3’-untranslated region (3’UTR) of specific target messenger RNAs (mRNAs), causing mRNA degradation or translational repression (1). A single miR can regulate numerous target mRNAs and conversely, a single mRNA can be targeted by several miRs. miRs have a potential role to play in tumor development and sustenance (oncomirs) by down-regulating tumor suppressor genes (2) but they can also act as tumor suppressors in a highly tissue-specific manner (3). The downregulation of tumor suppressor genes is essential for continuous proliferation and survival of cancer cells. One such important tumor suppressor protein is programmed cell death 4 (Pdcd4), which is downregulated in various cancers of oral (4), breast (5), lung (6), liver (7), brain (8) and colon (9) tissues. Pdcd4 plays an important role in regulating apoptosis, invasion, and metastasis (10-12) and is known to inhibit AP-1-dependent transcription (13, 14). miR-155 mediated downregulation of Pdcd4 was suggested earlier (15) and a recent study in lung cancer has claimed that microRNA-155 (miR-155) directly targets Pdcd4 and down-regulates its expression in lung cancer cells (16) but did not provide a convincing evidence for Pdcd4 3’UTR mediated downregulation as a mechanism. MicroRNA-155 (miR-155) was found to be overexpressed in oral (17), breast (18), tongue (19), pancreatic (20), prostatic (21) and thyroid cancers (22). miR-155 over expression has also been implicated in enhanced cell proliferation, invasion and metastasis(23) by downregulating the expression of tissue specific target genes such as APC (24), CDC73 (25), DET1 (26), FOXO3a (27), SMAD2 (28) and TP53INP1 (20) in many cancers. miR-155 is generated as a non-coding mRNA from B-cell integration cluster (BIC) gene located on chromosome 21 whose promoter is mainly controlled via Activation protein-1 (AP-1), (−36bp upstream of start site) and NF-κB (nuclear factor kappa-light-chain-enhancer of activated B cells), (−1150 upstream from start site) transcription factors present upstream of core promoter region (29). AP-1, formed by the heterodimerization between members of JUN and FOS protein families, is a critical regulator of cell proliferation, apoptosis, tumor invasiveness, angiogenesis and other multiple hallmarks of cancer (30-32).

In this study, we demonstrate that miR-155 is overexpressed in tongue cancer cells (SAS and AWL) and tongue tumor tissues when compared to normal FBM cells and normal tongue tissues, respectively. We hypothesized that miR-155-mediated targeting of 3’UTR of Pdcd4 mRNA results in the down-regulation of Pdcd4 in tongue cancer and this may account for the increased activation of AP-1-dependent transcription of BIC promoter which results in overexpression of miR-155. Here we demonstrate for the first time that miR-155 directly targets Pdcd4 in tongue cancer cells and indirectly activates AP-1-dependent transcription of BIC gene, and thus promotes overexpression of miR-155 by the miR-155/PDCD4/AP-1 positive feedback loop in tongue cancer

## 2 RESULTS

### 2.1 miR-155 expression shows inverse relation to that of its predicted target, Pdcd4, in tongue cancer cells and tissues

A panel of cancer cell lines of different tissue origin was screened for the endogenous expression of miR-155 and the results show that miR-155 is significantly higher in tongue cancer cells (SAS and AWL) than others tested (Figure. 1A). Next, we checked its expression in a panel of head and neck carcinoma cells and an immortalized fetal buccal mucosal cell line (FBM). We found that miR-155 expression was less in FBM and was relatively high in head and neck cancer cells with the highest levels being noticed in tongue cancer cells (SAS and AWL cells) (Figure. 1B) and interestingly Pdcd4 is poorly expressed in tongue cancers cells (SAS and AWL cells) (Figure. 1C). In-silico target prediction software’s predicted Pdcd4 as a target of miR-155, supported by the presence of miR-155 seed sequence match (100%) from nucleotide 1774 to 1783 in the 3’UTR of Pdcd4 with the free energy (dG) of −21.4Kcal/mol (Supplementary Figure. S1 B). All these result clearly indicates that SAS and AWL cells shows inverse relationship for the expression levels of miR155 and Pdcd4 (Figure. 1B and 1C) as often seen in miRNA-target pairs in cancer cells (33).

**Figure. 1.**
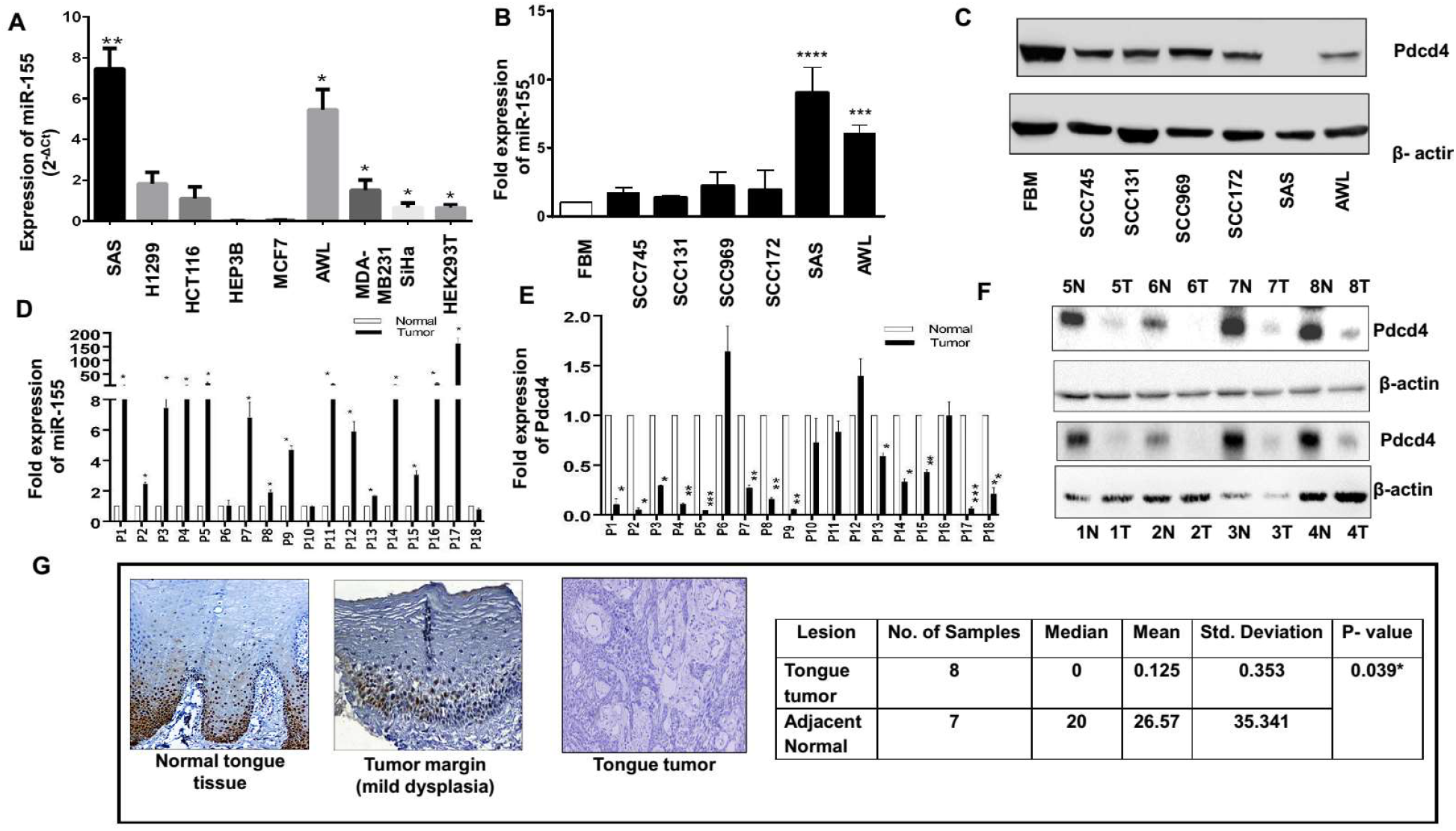
Analysis of miR-155 and Pdcd4 expression levels in cell lines and tongue tumors. **A**, Endogenous expression of miR-155 in cancer cell lines of different tissue origin from three different passage calculated by 2-ΔCt method (statistical analysis is performed column statistics with one-sample t-test n=3) **B,** Fold expression of miR-155 in FBM and different head and neck carcinoma cells normalized to U6 (n=3) **C,** Western blot for Pdcd4 and β-actin (as internal control gene) in FBM and different head and neck carcinoma cells. **D,** Fold expression of miR-155 normalized to U6 in normal and tumor tongue tissues (n=3 as experimental triplicates). **E,** Fold expression of Pdcd4 mRNA normalized to β-actin in normal and tumor tongue tissues (n=3 as experimental triplicates). **F,** Western blot for Pdcd4 and β-actin for the tongue cancer tissue samples with adjacent normal tissues - Normal (N) and Tumor (T) for patients P1 to P8. **G,** Representative Immunohistochemistry images for Pdcd4 in Normal tongue tissue, tumor margin (mild dysplasia) and tongue cancer tissue at 20X magnification. Values are expressed as the mean ± SD (*p < 0.05, **p < 0.01, ***p < 0.001).

Since miR-155 and Pdcd4 expression levels in SAS and AWL cells show inverse relation, it was relevant to check if there is an inverse relationship in their expression in tongue cancer tissues and to this end, we analyzed miR-155 and Pdcd4 levels in 18-pairs of tongue cancer patient samples (adjacent normal and tumor). miR-155 expression was found to be higher in most of the tumor tissues (except patient 6) compared to their adjacent normal tissue sections when analyzed by qPCR (Figure. 1D). As expected, Pdcd4 mRNA levels were lower in tongue cancer tissues in comparison to their adjacent normal tissues (Figure. 1E). The protein levels of Pdcd4 were detected by western blot for 8 patients and immunohistochemistry for rest for other 18 patients and the results show that Pdcd4 was barely detected in tumor samples compared to normal tissue (Figure. 1F and 1G). These data indicate the existence of inverse correlation in the expression levels of miR-155 and its target, Pdcd4 and prompted us to experimentally validate for the first time if Pdcd4 is a target of miR-155 in tongue cancer cells.

### 2.2 miR-155 negatively regulates Pdcd4 expression

First, we have cloned the wild-type 3’UTR of Pdcd4 (nucleotides 1170-1913) and miR-155 binding site mutant 3’UTR of Pdcd4 into psiCheck-2 (Supplementary Figure. S2 A). Since FBM and SCC745 cells express low and moderate levels of miR-155 (Figure. 1B), respectively, they were co-transfected with pcDNA-*BIC* expressing miR-155 and psiCheck-3’UTR of Pdcd4 WT or MUT. The normalized renilla luciferase activity of psiCheck-3’UTR WT was significantly reduced upon ectopic expression of miR-155 in both FBM (Figure. 2A) and SCC745 cells (Supplementary Figure. S2 B) but not much change in that of psiCheck-Pdcd4 3’UTR MUT (Figure. 2A and Supplementary Figure. S2B). Further, FBM and SCC745 cells were transfected with increasing amounts of pcDNA-*BIC* or pcDNA3.1 (+) (1µg to 4µg) as control and a gradual increase in the expression levels of miR-155 over the control were noticed using qPCR analysis (Figure. 2B and Supplementary Figure. S2C). When analyzed for the expression of Pdcd4 mRNA, there was a gradual decrease in the levels of Pdcd4 mRNA in FBM cells (Figure. 2C) and SCC745 (Supplementary Figure. S2D). The protein levels of Pdcd4 showed a gradual decrease with increase in the expression of miR-155 in FBM (Figure. 2D) and SCC745 cells (Supplementary Figure. S2E). These results suggest that overexpression of miR-155 has the potential to target Pdcd4 and downregulate its expression in FBM and SCC745 cells.

**Figure 2.**
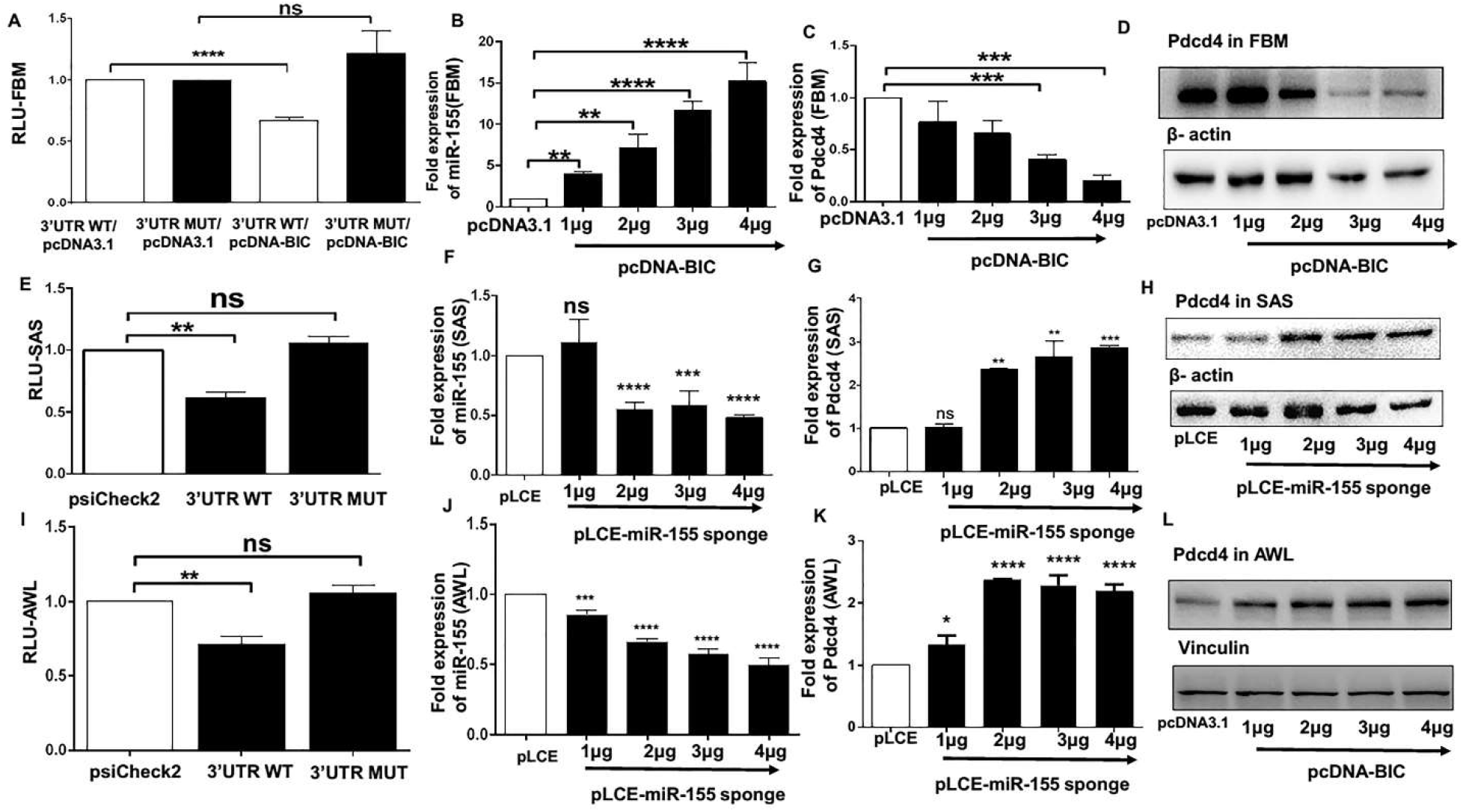
Targeting of 3’UTR of Pdcd4 by miR-155 and the consequent changes in the expression of Pdcd4 in FBM, SAS and AWL cells. Dual luciferase reporter assay was performed in **A,** FBM cells by cotransfecting them with WT or MUT (100ng) 3’UTR Pdcd4 and pcDNA3.1 or pcDNA-BIC and the graphs represent normalized values (RLU-FBM) of renilla/firefly luciferase activities (n=3) **B,** Fold expression of miR-155 normalized to U6 in FBM cells transfected with different amounts (1µg to 4µg) of pcDNA-BIC with pcDNA3.1 as a control (n=3). **C,** Fold expression of Pdcd4 mRNA in FBM cells transfected with different amounts (1µg to 4µg) of pcDNA-BIC with pcDNA3.1 as a control normalized to β-actin mRNA (n=3). **D,** Western blot for Pdcd4 and β-actin in FBM cells ransfected with different amounts (1µg to 4µg) of pcDNA-BIC with pcDNA3.1 as a control. **E,I,** Dual luciferase reporter assay performed in SAS and AWL cells transfected with 100ng of psiCheck-2, psiCheck-Pdcd4 3’UTR WT or psiCheck-Pdcd4 3’UTR MUT (n=3). **F, J,** Fold expression of miR-155 in SAS and AWL cells transfected with pLCE-miR-155 sponge plasmid from 1 to 4µg with 4µg pLCE as a control (n=3). **G, K,** Fold expression of Pdcd4 mRNA in SAS and AWL cells transfected with pLCE-miR-155 sponge plasmid from 1 to 4µg with 4µg pLCE as a control,normalized to β-actin (n=3). **H, L,** Western blot for Pdcd4 in SAS and AWL cells when transfected 1 µg to 4µg pLCE-miR-155 sponge plasmid with 4µg pLCE as control,β-actin and vinculin are taken as internal control respectively. Values are expressed as the mean ± SD (*p < 0.05, **p < 0.01, ***p < 0.001).

As SAS and AWL cells express high levels of miR-155 (Figure. 1A and 1B), they were transfected with psiCheck-2, psiCheck-3’UTR of Pdcd4 WT or MUT, and as expected, the renilla luciferase activity showed a significant decrease with psiCheck-Pdcd4 WT but not the other two constructs (Figure. 2E and 2I). Next, we decreased the expression of miR-155 in SAS and AWL cells using miR-155 sponge constructs containing 9 tandem repeats of miR-155 binding sites. Ectopic expression of pLCE-miR155 sponge (1 to 4µg) with pLCE as control resulted in a decrease in endogenous levels of miR-155 (Figure. 2F and 2J) and Pdcd4 levels increased both at the mRNA (Figure. 2G and 2K) and protein levels (Figure. 2H and 2L). Taken together, these results show that miR-155 regulates Pdcd4 expression by binding to its 3’UTR in FBM, SCC745, SAS and AWL cell lines.

### 2.3 Feedback loop between miR-155, Pdcd4 and AP-1 provides a mechanistic basis for the overexpression of miR-155 in tongue cancer

To understand whether *BIC* promoter regulation involves miR-155, Pdcd4 and AP-1, HEK293T and SAS cells were transfected with pGL-3 *BIC* promoter and pcDNA3.1 (-) or pcDNA-Pdcd4. The luciferase activity of pGL3-*BIC* promoter was markedly reduced when co-transfected with pcDNA-Pdcd4 compared to pcDNA3.1 (-) (HEK293T-supplementary Figure. 3A and SAS-Figure. 3A). To validate involvement of AP-1 or NF-κB transcription factors in *BIC* gene promoter regulation in these cells, we used promoter constructs mutated at the AP-1 or NF-κB binding sites together with pcDNA3.1 (-) and pcDNA-Pdcd4. We observed that mutation of the conserved AP-1 binding site reduced *BIC* promoter activity, whereas mutation at the NF-κB binding site had less effect in both cell lines transfected with pcDNA3.1 (-) (HEK293T-supplementary Figure. 3A and SAS-Figure. 3A). However, in pcDNA-Pdcd4 transfected cells, the effect was more significant in reducing the *BIC* promoter activity in the case of NF-κB mutant (with wt AP-1) but not in the AP-1 mutant (with wt NF-κB) (HEK293T-supplementary Figure. 3A) and (SAS-Figure. 3A). Therefore, we conclude that Pdcd4 regulates *BIC* promoter primarily by AP-1 driven transcription in HEK293T (supplementary Figure. 3A) and SAS cells (Figure. 3A).

**Figure. 3.**
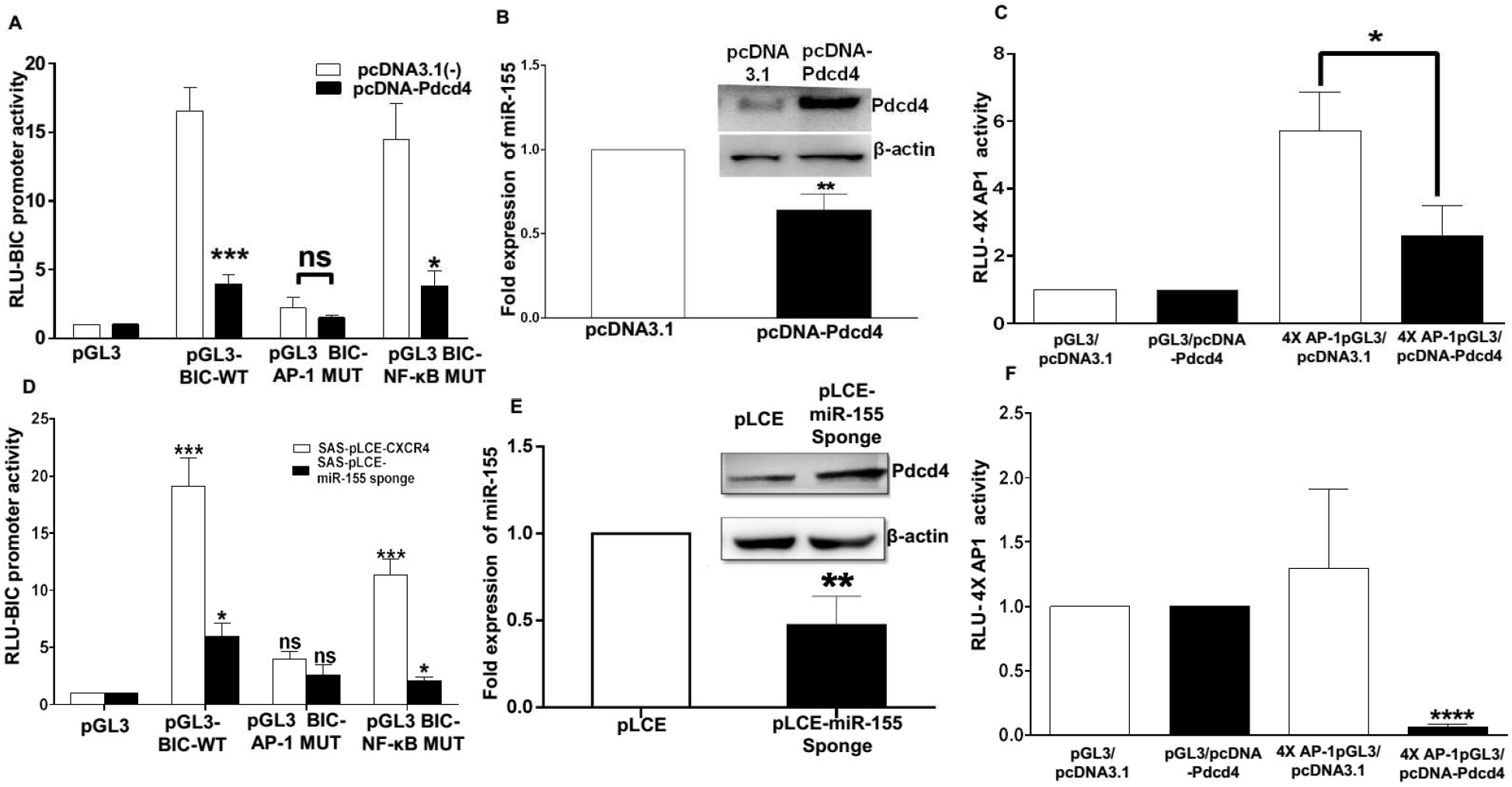
Feedback loop between miR-155, Pdcd4 and AP-1 in SAS cells. **A,** Dual luciferase reporter activity of wild type BIC promoter and BIC promoter mutated at AP-1 or NF-κB binding site in presence of pcDNA3.1(-) or pcDNA-Pdcd4 in SAS cells (n=3). **B,** Fold expression of miR-155 normalized to U6 upon over expression of Pdcd4 in SAS cells (n=3). Insets show western blotting of Pdcd4 and β-actin in SAS cells in presence of pcDNA3.1 (-) or pcDNA-Pdcd4. **C,** AP-1 luciferase activity detected by 4X-AP-1 binding sites cloned upstream of pGL3 luciferase plasmid in presence of pcDNA-Pdcd4 and pcDNA3.1(-) in SAS cells (n=3). **D,** Luciferase activity of wild type BIC promoter and BIC promoter separately mutated at AP-1 or NF-κB binding site in presence of pLCE and pLCE-miR-155 sponge in SAS cells (n=3). **E,** Fold expression of miR-155 normalized to U6 upon ectopic expression of pLCE and pLCE-miR-155 sponge in SAS cells (n=3). (Inset) Western blots for expression of Pdcd4 and β-actin in SAS cells upon ectopic expression of pLCE and pLCE-miR-155 sponge. **F,** AP-1 activity detected by 4X-AP-1 luciferase construct in presence of pLCE and pLCE-miR-155 sponge in SAS cells. Values are expressed as the mean± SD (*p < 0.05, **p < 0.01, ***p < 0.001).

Upon ectopic expression of Pdcd4 (inset Figure. 3B) miR-155 levels reduced in HEK293T (supplementary Figure. 3B) and SAS cells (Figure. 3B). To validate the regulation of AP-1 transcription by Pdcd4, we have cotransfected HEK293T and SAS cells with pGL3 basic or 4X-AP-1 luciferase construct with pcDNA-Pdcd4 or pcDNA3.1 (-) and this decreased the AP-1 luciferase activity over that of corresponding controls in presence of Pdcd4 in both HEK293T (supplementary Figure. 3C) and SAS cells (Figure. 3C).

To gain further insights into the regulation of *BIC* promoter by feedback loop between miR-155/Pdcd4/AP-1, we have cotransfected HEK293T with pGL3-*BIC* and pcDNA-*BIC* or pcDNA3.1 (+) and we have found significant increase in luciferase activity of *BIC* promoter over the corresponding controls, while BIC-AP-1 mutant is not much affected in the presence of miR-155 (Supplementary Figure. 3D). The *BIC*-NF-kB mutant shows reduced activity in presence of miR-155 indicating the involvement of AP-1 site which is in native form in this construct (Supplementary Figure. 3D). Overexpression of miR-155 was confirmed by qPCR (Supplementary Figure. 3E) and Pdcd4 levels were quantified by western blot (Inset). To validate the regulation of AP-1 transcription by miR-155, we have cotransfected HEK293T cells with pGL3 basic or 4X-AP-1 luciferase construct with pcDNA-*BIC* or pcDNA3.1 (+) and this increased the AP-1 luciferase activity over that of corresponding controls in presence of miR-155 in both HEK293T cells (Supplementary Figure. 3F).

As SAS cells have higher expression of miR-155, we have cotransfected them with the pGL3-*BIC* and pLCE or pLCE-miR-155 sponge and it was found that *BIC* promoter activity was reduced in presence of miR-155 sponge which results in higher expression of Pdcd4 (Figure. 3D). The reduced expression of miR-155 expression is confirmed by miR-155 specific stem-loop qPCR (Figure. 3E). The upregulation of Pdcd4 was confirmed by western blotting, on ectopic expression of miR-155 sponge (Figure. 3E {inset}). The decreased AP-1 luciferase activity is seen upon cotransfection of 4X-AP-1 luciferase construct with a pCLE-miR-155 sponge when compared with pCLE (Figure. 3F). All these experimental pieces of evidence indicate the existence of feedback loop operating in SAS cells which may be responsible for high expression of miR-155 in these cells.

### 2.4 miR-155 sponge depletes miR-155 and halts tumorigenic properties of SAS cells

For gaining more mechanistic insights, we generated stable clones of SAS cells that express miR-155 sponge using lentiviruses. The virus titer was optimized for efficient transduction and positively transduced cells were enriched by cell sorting in FACS-ARIA III using GFP as a marker (Supplementary Figure. 4A). Real-time PCR for miR-155 shows significant down-regulation of miR-155 expression in pLCE-miR-155 sponge SAS compared to pLCE-SAS cells (Figure. 4A). Pdcd4 is upregulated both at transcriptional and translational levels in pLCE-miR-155 sponge SAS compared to pLCE-SAS cells (Supplementary Figure. 4B and Figure. 4B). To further confirm the effectiveness of sponge, we have used miR-155 luciferase-reporter sensor system having a 3X exact binding sequence for miR-155 and found that the luciferase activity is almost 7-fold high in pLCE-miR-155 sponge stable cells as compared to pLCE-SAS cells (Figure. 4D). Furthermore, pLCE-SAS cells and pLCE-miR-155 sponge SAS cells were transfected with psiCheck-2, psiCheck-3’UTR of Pdcd4 WT or MUT. As expected, the renilla luciferase activity shows a significant decrease with psiCheck-Pdcd4 WT in pLCE-SAS cells but not in pLCE-miR-155 sponge SAS cells and the psiCheck-3’UTR of Pdcd4 MUT (Figure. 4C). All these experimental evidence clearly show that miR-155 directly targets Pdcd4 in SAS cells and depletion of miR-155 upregulates the expression of Pdcd4.

**Figure. 4.**
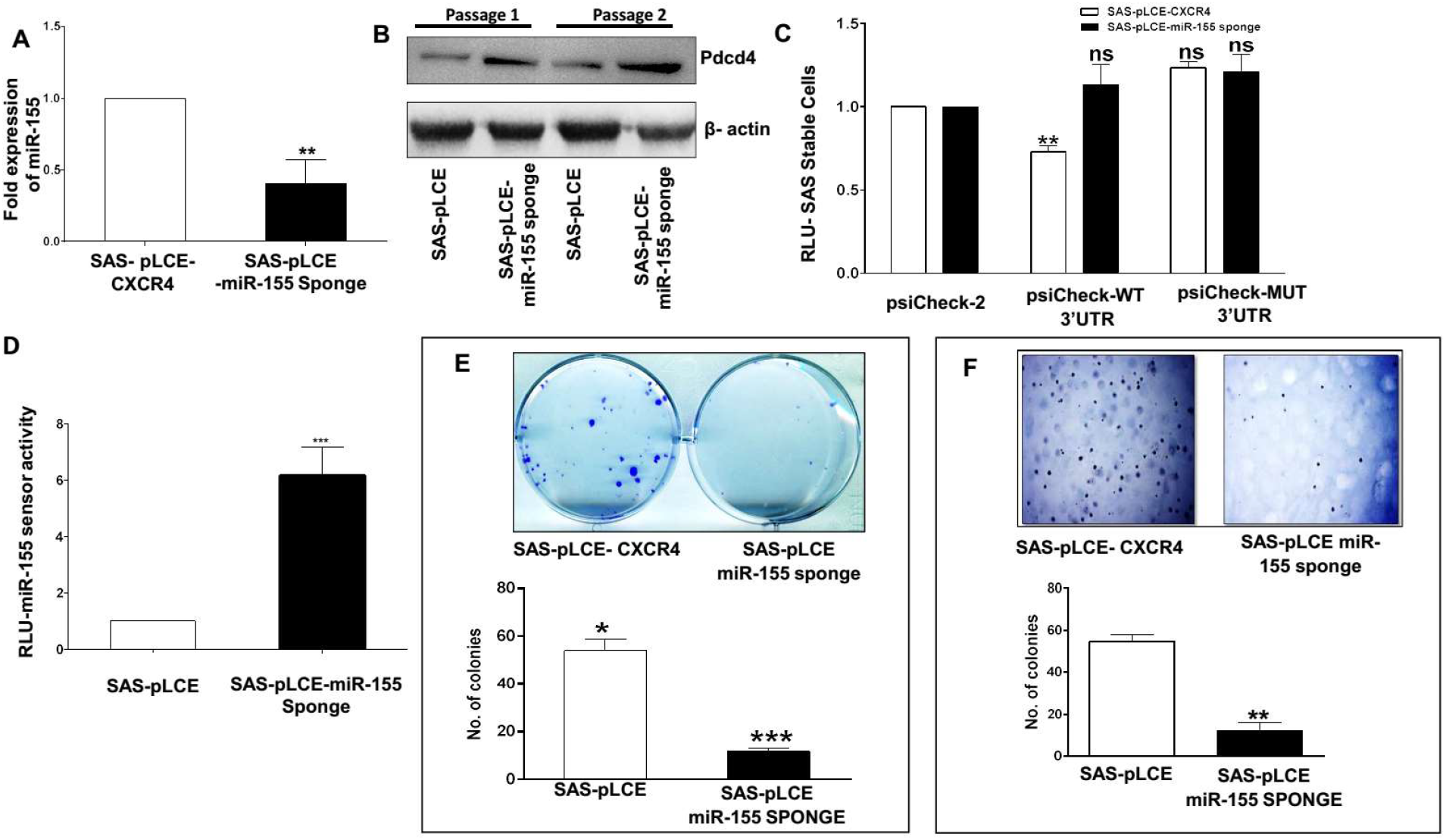
Lentiviral based stable expression of miR-155 sponge and the consequent changes in SAS cells. **A,** Fold expression of miR-155 between SAS-pLCE and SAS-pLCE-miR-155 sponge normalized to U6 (n=3). **B,** Western blot of Pdcd4 and β-actin for SAS-pLCE and SAS-pLCE-miR-155 sponge with two different cell passages **C,** Dual luciferase assay was performed in SAS-pLCE and SAS-pLCE-miR-155 sponge cells transfected with 100ng of psiCheck-2, psiCheck-Pdcd4 3’UTR WT or psiCheck-Pdcd4 3’UTR MUT (n=3). **D,** Dual luciferase reporter assay performed with pGL-3X complementary binding sites for miR-155 /miR-155 sensor plasmid (n=3). **E,** Representative images of clonogenic assay with SAS-pLCE and SAS-pLCE-miR-155 sponge cells and the graph shows the mean number of colonies from three separate experiments (n=3). **F,** Representative images of soft agar assay with SAS-pLCE and SAS-pLCE-miR-155 sponge cells and the graph is a representation of number of colonies from three separate experiments (n=3). Values are expressed as the mean ± SD (*p < 0.05, **p < 0.01, ***p < 0.001).

Clonogenic or colony formation assay is an in-vitro cell survival assay designed to measure the ability of a single cell to grow into a colony(34). There was a drastic decrease in the ability of SAS-pLCE-miR-155 sponge single cells to grow into colonies compared to SAS-pLCE-stable SAS (Figure. 4D). Also, tumour cells have the propensity to grow in an anchorage-independent manner and hence can form colonies in a semi-solid medium such as soft agar (35). The anchorage-independent growth of the stable pLCE and pLCE-miR-155 sponge cells was analyzed by the soft agar assay in 6 well tissue culture plates for 24 days. The colony formation capability on soft agar decreases in pLCE-miR-155 sponge cells compared to pLCE-SAS cells (Figure. 4E). These results suggest that reducing the miR-155 levels in SAS cells reduces their survival and ability to form colonies on soft agar.

### 2.5 miR-155 sponge inhibits cell viability, arrests cell cycle and induces apoptosis in SAS cells

Since Pdcd4 is known to be upregulated upon induction of apoptosis (36), it was of interest to check the changes in cellular apoptosis, cell cycle and cell viability. First, we monitored cell viability at different time points by MTT assay and found SAS-pLCE-miR-155 sponge cells exhibited a gradual decrease in viability with increase in time but not the SAS-pLCE cells (Figure. 5A). Next, cell cycle analysis (by propidium iodide) of SAS-pLCE-miR-155 sponge cells shows an increase in a number of cells in sub-G0 and G2/M phase of the cell cycle when compared with SAS-pLCE cells (Figure. 5B). The increased percentage of annexin V stained cells were found upon down-regulation of miR-155 in SAS cells (Figure. 5C). As Pdcd4 is among first few proteins upregulated during the process if apoptosis, it was of interest to analyze the important pro-apoptotic proteins like cleaved caspase 3, cleaved caspase 9 and cleaved PARP and as expected we found them upregulated in SAS-pLCE-miR-155 sponge transfected cells compared to SAS-pLCE cells (Figure. 5D). We also analyzed the expression of Bcl2 and Bax and found that Bax but not Bcl2 was upregulated in SAS-pLCE-miR-155 sponge cells compared to SAS-pLCE cells (Figure. 5E). Taking together these results show cell cycle arrest at sub-G0 and G2/M phase together with an increased number of annexin V positive in SAS-pLCE-miR-155 sponge compared to SAS-pLCE cells. Also increased expression of cleaved caspase 3, cleaved caspase 9, cleaved PARP, Bax indicate that miR-155 depletion in SAS cells leads to a decrease in cell viability and the induction of cell cycle arrest and apoptosis.

**Figure. 5.**
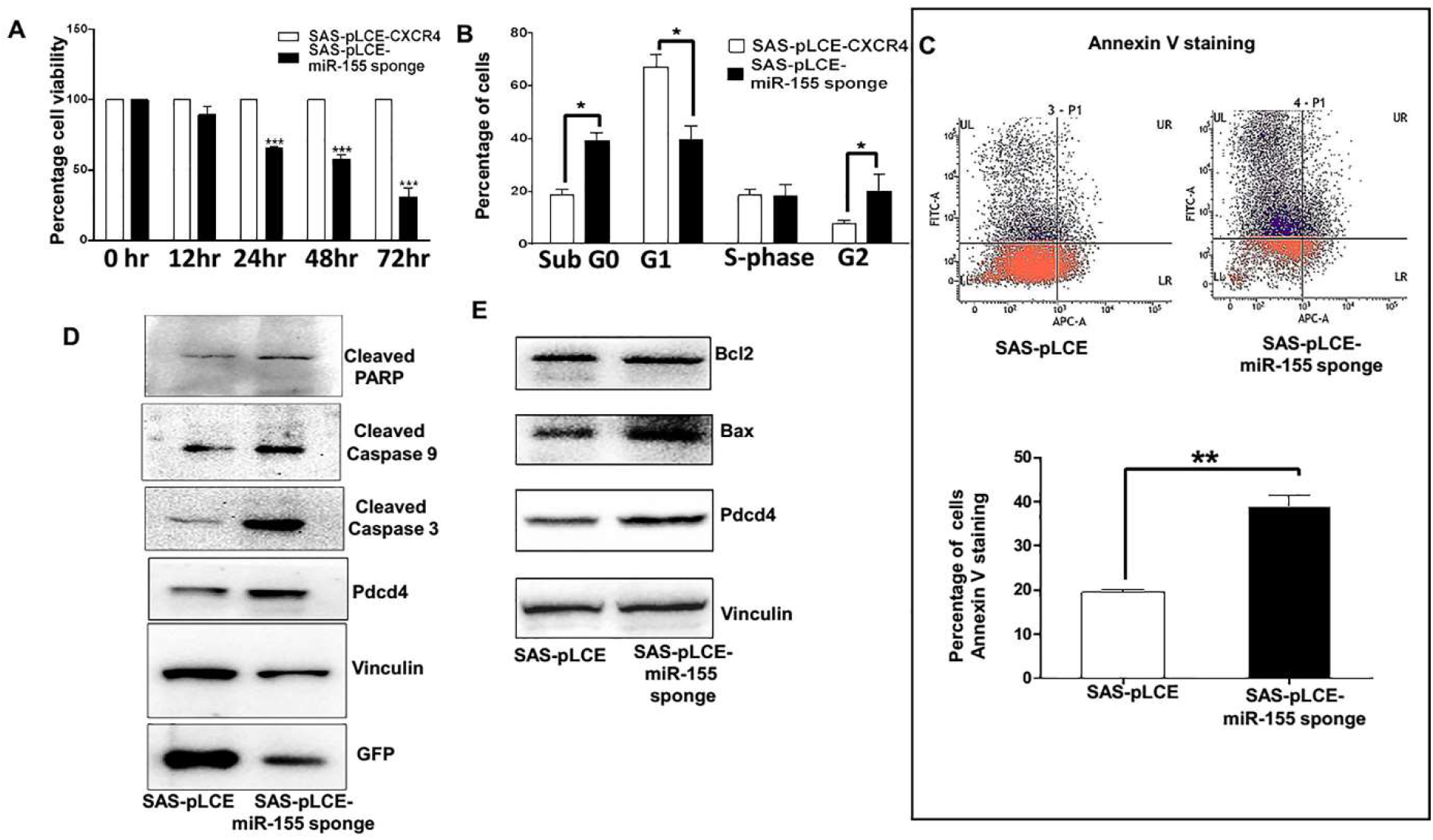
Effect of miR-155 inhibition on cell viability, cell cycle and apoptosis. **A,** Effect of miR-155 depletion on cell viability as measured by MTT assay (n=3). **B,** Cell cycle analysis by propidium iodide of SAS-pLCE and SAS-pLCE-miR-155 sponge cells, each bar in the graph represent percentage of cells present in particular phase of cell cycle (n=3). **C,** Effect of miR-155 down regulation on cellular apoptosis detected by Annexin V staining (n=3). **D,** Western blot for cleaved parp, cleaved caspase 9, cleaved caspase 3, Pdcd4 and vinculin (as internal control). GFP levels acts as an indirect indicator that miR-155 is effectively inhibited by miR-155 sponge. **E,** Western blot for Bcl2, Bax, Pdcd4 and Vinculin (acting as internal control). Values are expressed as the mean ± SD (*p < 0.05, **p < 0.01, ***p < 0.001).

### 3.6 miR-155 knockdown reduces xenograft formation in nude mice

Consistent with the above in-vitro results, nude mice injected with 5×10^6^ SAS pLCE-miR-155 sponge cells showed a significant reduction in tumor burden over SAS-pLCE cells. The growth kinetics recorded at 6 different time points post-injection and the tumor volumes (in mm3) measured are shown in (Figure. 6A). Tumor weights were also measured at the end of the study, the weight for the xenografts generated by SAS pLCE-miR-155 sponge cells was significantly lower compared to xenografts generated by SAS-pLCE-cells (Figure. 6B). The expression of miR-155 is elucidated by qPCR from the RNA collected from tumor xenografts generated by SAS pLCE-miR-155 sponge and SAS pLCE cells. We have also performed western blot for Pdcd4 for the same set of tumors and it was found that SAS-miR-155 sponge cells show higher expression of Pdcd4 protein compared to SAS-pLCE cells (Figure. 6C). We have also performed western blots for GFP from lysates of SAS-pLCE and SAS-pLCE-miR-155 sponge xenografts to clarify that tumors generated in mice are generated by injected cells (Figure. 6D). The in-vivo study clearly demonstrated that miR-155 knockdown suppressed tumor formation of SAS cells in nude mice.

**Figure. 6.**
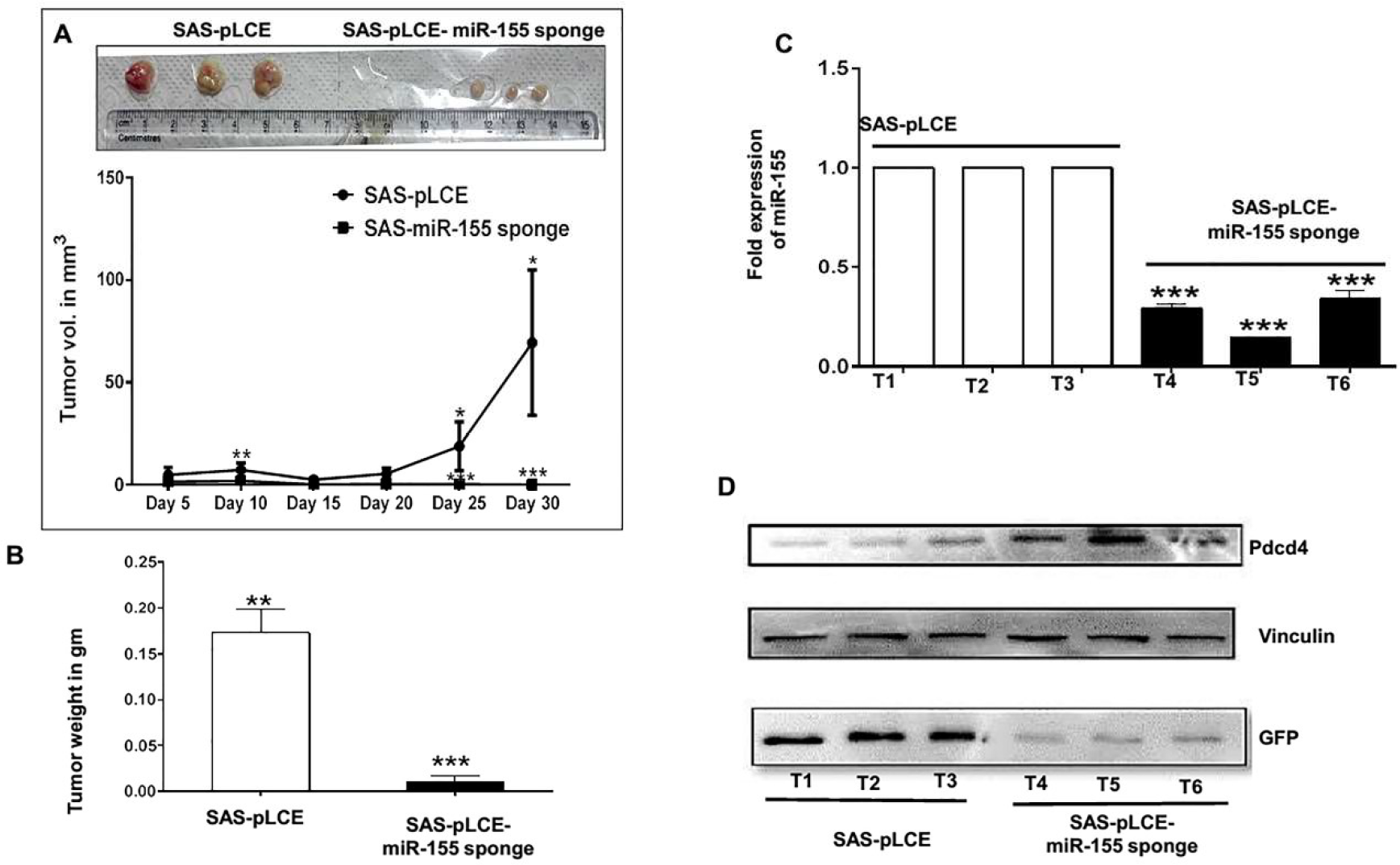
Effect of miR-155 sponge on tumor growth in nude mice. **A,** Representative images showing the tumor formation in nude mice with graph showing the changes in tumor volume (n=3). **B,** Changes in the weight of tumor xenografts produced by SAS-pLCE-miR-155 sponge and SAS-pLCE cells (n=3) **C,** Fold expression of miR-155 in the tumors by qPCR (n=3 as experiemtal triplicates) and **D,** Western blots for Pdcd4 with vinculin acting as internal control, GFP act as indicator of effective inhibition of miR-155 in xenografts. T1, T2 and T3 represent xenografts from mice injected with SAS-pLCE cells whereas T4, T5 and T6 represent xenografts from mice injected with SAS-pLCE-miR-155 sponge cells. Values are expressed as the mean ± SD (*p < 0.05, **p < 0.01, ***p < 0.001).

## 3 Materials and methods

### 3.1 Cell lines, reagents and plasmids

The cell lines, Hep3B, SiHa, MCF7, MDA-MB231 and H1299, were obtained from NCCS, Pune, India. HCT116 and HCT116P53-/-cells were received as a gift from Bert Vogelstein, Johns Hopkins University School of Medicine, USA. SCC131, SCC745, SCC969, AWL and SCC172 cell lines were obtained as a gift from Suresh Kumar Rayala, IIT Madras. SAS (Tongue carcinoma cell line) was procured from the Japanese Collection of Research Bioresources Cell Bank, Japan. Cell lines were maintained in DMEM with 10% FBS (Invitrogen) containing penicillin and streptomycin as antibiotics. FBM cell line was received as gift from Milind Vaidya (ACTREC) Tata Memorial Centre, India and maintained in IMDM + 10%FBS supplemented with insulin, Hydrocortisone and EGF. The pcDNA-*BIC* plasmid was a gift from Dr Iqbal Rather, IISc, Bangalore. The pLCE and pLCE-miR-155 sponge were gifted by Bryan R. Cullen, Dept. of Molecular Genetics & Microbiology, Duke University, UK. pPax2 and pMDL-2 were obtained from Addgene, USA. All the pGL-3 *BIC* promoter plasmids with their respective AP-1 and NF-κB mutant plasmids was a kind gift from Erik K. Flemington, Tulane University, USA.

### 3.2 Construction of 3’UTR reporter for Pdcd4

The 688bp of 3’UTR of PDCD4 was amplified using gene-specific primers (Forward Primer - 5’ATTACCTAACGTGACATGGCACATAAAATTGGTTAAAAAATTTTG3’ and Reverse primer 5’-TAATGGATATCTTGTGTGACCAGATCCCACCAGTAATG -3) containing restriction sites for XhoI and NotI to facilitate the directional cloning of 3’UTR of Pdcd4 in a psiCheck-2 plasmid. The 3’UTR for Pdcd4 was mutated at miR-155 seed recognition region by overlap extension PCR. The insert sequences for both wild-type and mutated 3’UTR of Pdcd4 were verified by DNA sequencing performed by Eurofins Genomics, India.

### 3.3 3’UTR luciferase assay

FBM and SCC745 were cotransfected with pcDNA3.1 or pcDNA3.1-*BIC* and the wildtype or mutant 3’UTR luciferase, whereas AWL and SAS cells were transfected only with psiCheck-2, 3’UTR WT, 3’UTR MUT constructs and after 48 h post transfection, cells were lysed using Passive Lysis Buffer, and Renilla luciferase activity was measured using the Dual Luciferase Assay Kit (no. A2492, Promega) and a luminescence plate reader (Molecular Devices Inc., Sunnyvale, CA, USA), wherein firefly luciferase acted as the internal control.

### 3.4 BIC and AP-1 Promoter luciferase assay

The regulation of *BIC* and 4X-AP-1 promoter by Pdcd4 or miR-155 was performed using different combination of plasmid constructs in HEK293T and SAS cells Table S1, Table S2, Table S3 and Table S4. Further downstream process was performed after 48 h post transfections, by lysing the cells in Passive Lysis Buffer, and firefly luciferase activity was measured using the Dual Luciferase Assay Kit (no. A2492, Promega) in a luminescence plate reader (Molecular Devices Inc., Sunnyvale, CA, USA), wherein renilla firefly luciferase activity acted as the internal control.

### 3.5 Western blot and immunohistochemistry

All the cell lysates were prepared with RIPA cell lysis buffer and 30µg of protein lysates were loaded per well of 12% SAS-PAGE gel and later transferred to PVDF membrane at a constant current of 220mA for 1-1/2 hours. The membrane was probed by the primary antibody, washed thrice with TBS-T and TBS separately and again probed with secondary antibody conjugated with HRP. All western blots were developed in ChemiDoc western blot apparatus from BioRad. For IHC, the tissue slide was incubated with 1:500 dilution of primary antibody. The slides were scored by the pathologist as the percentage positivity of nuclear or cytoplasmic staining and were graded as a) negative<=5%; b) weak, 5-19%; c) moderate, 20-49%; and d) strong, 50-100%) [26].

### 3.6 Transfection, RNA extraction, cDNA synthesis and Real-time PCR

We used linear chain PEI for plasmid DNA transfections unless mentioned otherwise [27]. We have optimized transfection efficiency at 2:1 PEI: DNA for FBM, HEK293T, SAS, AWL and SCC745 cells. The tissue samples were stored in RNase later and stored at −80°C until further use. Total RNA from cells was collected by a TriZol reagent (Invitrogen) as per the manufacturer’s protocol. For miRNA cDNA synthesis, stem-loop primer-based PCR protocol was used [28]. For cDNA synthesis, oligodT based conversion of mRNA to cDNA is performed by MMLV reverse transcriptase from life technologies. miRNA-155 expression was quantified by performing stem-loop reverse transcription followed by quantitative PCR; reverse transcription by MMLV reverse transcriptase (Life Technologies) was performed using miR-155 -specific and U6-specific stem-loop primers. qRT-PCR analysis of miR-155 and Pdcd4 is performed by using sybr green chemistry and quantified by 2-ΔCt and 2-ΔΔCt [29]. The expression of miR-155 is normalized to U6 snRNA and Pdcd4 to β-actin. All qRT-PCR reactions were performed in triplicate.

### 3.7 Production of lentiviruses for stable expression of miR-155 sponge

Lentiviruses were produced by co-transfecting HEK293T cells with pPax2, pMDL-2 and pLCE-miR-155 sponge/pLCE. SAS cells were transduced with the optimized virus titer and incubated in presence of 8 µg/ml polybrene. The positively transduced cells were enriched by cell sorting in FACS-ARIA III by using eGFP to sort the positive clones.

cells were enriched by cell sorting in FACS-ARIA III by using eGFP to sort the positive clones.

### 3.8 Soft agar assay

The cancerous cells have the tendency to grow in an anchorage-independent manner and hence have a tendency to form colonies in a semi-solid medium such as soft agar. The anchorage-independent growth of the SAS-pLCE and SAS-miR-155 sponge cells was analyzed by the soft agar assay in 6 well tissue culture plates. DMEM 10% FBS with 0.7% soft agar is poured first in each well of 6 well plate with utmost precision not to include air bubbles, later 3000 cells were mixed with 1 ml of DMEM 10% FBS with 0.35% soft agar and poured over base agar. The cells were allowed to grow in 5% C02 incubator at 37°C for 25 days with the addition of 500µl for liquid DMEM containing 10% FBS after every three days.

### 3.9 Clonogenic assay

For the clonogenic assay, 300 SAS-pLCE and SAS-miR-155 sponge cells were plated per well in 6 well plates. Cells were allowed to form colonies for 25 days and later fixed in methanol and stained with crystal violet. A colony of cells was considered as prominent if it contained 50 cells or more.

### 3.10 Cell cycle analysis

The cell cycle analysis of SAS-pLCE and SAS-miR-155 sponge cells was performed by the FACS-Calibur flow cytometer after staining the cells with propidium iodide. We have followed the end to end protocol as found at (http://www.abcam.com/protocols/flow-cytometric-analysis-of-cell-cycle-with-propidium-iodide-dna-staining)

### 3.11 Cell viability assay

SAS cells were first transfected with pLCE and pLCE-miR-155 sponge plasmids in 6 well plate. After 24hrs of transfection 2000 cells were plated in each well of 96 well plate. The MTT reagent, a tetrazole, is added at 24hr time intervals (0 to 72 hr) which was reduced to formazan in the mitochondria of living cells. The absorbance of this coloured solution was quantified by measuring the absorbance at 570nm in a spectrophotometer

### 3.12 Annexin V-APC apoptosis assay

After transduction with lentivirus expressing miR-155 sponge and control, apoptosis was measured by Annexin V-APC. Cells subjected were cultured for another 72 hrs and then detached by 0.25% trypsin, washed twice with PBS, and stained by annexin V-APC for 10 min at room temperature according to the manufacturer’s instructions (BD Biosciences). Subsequently, flow cytometric analysis was performed with annexin V-APC staining.

### 3.13 In-vivo assay for tumor formation

To see the effect of restoration of Pdcd4 expression by inhibition of miR-155 on tumor growth, 5X106 SAS-pLCE or SAS-pLCE-miR-155 sponge cells resuspended in PBS with 50% matrigel matrix, were injected subcutaneously into the posterior flank of each female BALB/c athymic 6-week-old nude mouse. Tumor growth was monitored, and its volume was measured every 5 days till 35th day. Tumor volume (V) was calculated as L*W2/2, where L and W represent large and small diameters of tumor formed, respectively. Tumor weight was measured at the end of the study. All nude mice experiments were approved by the institutional animal ethics committee.

### 3.14 Statistical analysis

Experiments were carried out in triplicate. Data are presented in means plus or minus the standard deviation of the mean. The differences between test and control groups were analyzed using the Graph Prism Program. The statistical analysis is performed by column comparing between samples of different nature. Unpaired t-test is performed when comparing two samples involved in experiment. One way ANOVA with multiple comparison is performed when comparing the three or more samples involved in a single experiment. After performing statistical analysis, P-values were calculated and represented as *P< 0.05,**P< 0.01, ***P< 0.001 or ****P<0.0001.

## 4 DISCUSSION

The deregulation of gene expression is often associated with various cancerous phenotypes like proliferation, invasion, metastasis and migration. The identification of molecular players that play key roles in gene expression such as miRNAs are relevant for effective therapeutic strategies. Various experimental evidences indicate that some miRNAs like miR-21 (37) and miR-139 (38) and miR-139 [35] form feedback regulatory loops with their targets for their continuous expression to down-regulation their target tumor suppressor mRNA. In this study we have investigated that miR-155 is upregulated in tongue cancer and verified Pdcd4 as its potential target that generates an positive feedback loop with AP-1 to maintain the tumorigenic growth in tongue cancer cells. Even though Pdcd4 is recently shown to be a direct target of miR-155 in non-small cell lung cancer (16) but the data represented in the study is not convincingly shows that 3’UTR of Pdcd4 is a direct of miR-155, as miR-155 binding site mutant seems to be responding to presence of miR-155 and study title seems quite opposite to the study conducted and data represented. While as our data provide strong evidence that overexpression of miR-155 in tongue cancer cells is due to the positive feedback loop between miR-155/Pdcd4/AP-1. This is the first study to show that miR-155 modulates its own expression by down-regulation of Pdcd4 and activation of AP-1 dependent transcription of *BIC* promoter in the context of tongue cancer cells (Figure. 3). Through a series of in silico, in vitro, and in vivo approaches, we found that miR-155 plays a key role in regulating Pdcd4 by directly targeting its 3’UTR in tongue cancer cells and down-regulating its expression. miR-155 expression is upregulated in solid tumors of diverse origin (23, 39) and consistent with this, we demonstrated that miR-155 expression is significantly higher in tongue cancer tissues and SAS and AWL cells than in adjacent normal tissues and normal FBM cells, respectively, (Figure. 1), suggesting that high miR-155 expression is tightly associated with tongue cancer development. A significant negative correlation between miR-155 expression and that of Pdcd4 protein in maximum number of tongue tumor tissues and SAS and AWL cells observed by us supports the notion that miR-155-mediated control of Pdcd4 is operational in tongue cancer. We have provided several lines of evidence to show that Pdcd4 is directly targeted by miR-155 in FBM, SCC745, SAS and AWL cells for the first time and effect of miR-155 down-regulation on cancerous properties of tongue cancer cells was reflected in-vitro with a decrease in clonogenic capability and anchorage-independent growth on soft agar (Figure. 4D and 4E). Furthermore, restoration of Pdcd4 by knockdown of miR-155 was accompanied by decreased cell viability, enhanced apoptosis, and reduction of tumor growth in nude mice. miRNAs usually regulate a large set of targets and our data do not rule out the idea that there may be more targets for miR-155 in addition to Pdcd4. Indeed Pdcd4 has been shown to be targeted by miR-21 in various cancer like hepatocellular carcinoma(40), oral cancer(4), breast cancer (41), colon carcinoma (42)), miR-96 and miR-183 in glioma (43, 44), and miR-499 in oropharyngeal cancer (45).

With respect to the mechanism, our data show that miR-155 acts as a transcriptional regulator of the AP-1 complex by directly targeting Pdcd4, an inhibitor of AP-1 transcription. We found that AP-1 is critical for *BIC* promoter activity in tongue cancer cells whereas the contribution of NF-κB, was relatively minor. AP-1 activation is often paradoxical as it promotes or inhibits apoptosis in different cellular systems, but our data suggest its role in ensuring tongue cancer cell proliferation emphasizing the need to understand the cellular and extracellular context within which it functions (46). The inhibition of Pdcd4 by miR-155, in turn, contributes to the increase in AP-1 activity of *BIC* promoter that explains the enhanced expression of miR-155 in tongue cancer cells, thus revealing a novel miR-155/Pdcd4/AP-1 feedback regulatory mechanism. Inhibition of AP-1 transcription factor by pdcd4 involves Jun-Jun homodimers or Jun-Fos heterodimers(32). Taken together, our study validates tumour suppressor Pdcd4 as a target of oncogenic miR-155 and the positive feedback loop of miR-155-Pdcd4-AP-1 maintains the miR-155 mediated biological effects of the tongue cancer phenotypic effects (Figure. 7). Our data suggest that molecular strategies manipulate miR-155 expression and/or miR-155-Pdcd4-AP-1 feedback loop provide novel avenues to explore in tongue cancer therapeutic.

**Figure. 7.**
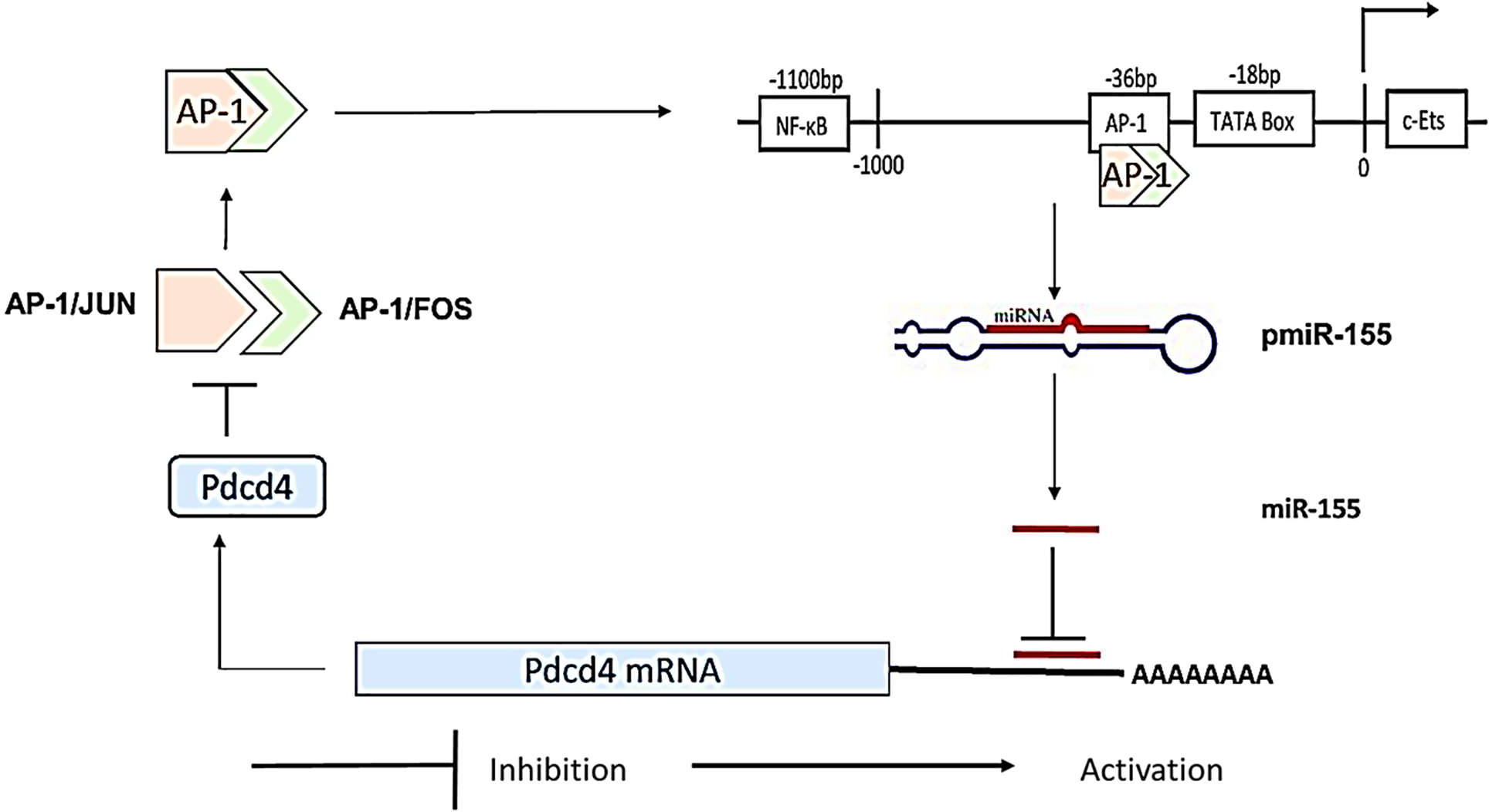
Overview of auto-regulatory feedback loop between miR-155/Pdcd4/AP1.

## 5 ACKNOWLEDGEMENTS

We thank DBT, Government of India, for the financial support (Grant No: BT/PR/2672/ AGR/36/ 702/ 2011), and Indian Institute of Technology Madras (IITM) for all other facilities. We are thankful to IIT Madras Alumni association for providing funds for travels to perform nude mice experiments. We would like to thank Dr Anuj Kaushik for his suggestions while writing this manuscript.

## 6 AUTHOR’S CONTRIBUTIONS

SZ performed in-vitro and in-vivo experiments, VT assisted in nude mice experiments under the supervision of KS, VS carried out the IHC experiments and scoring under the supervision of VR. SZ and DK analyzed the data. SZ and DK designed the experiments and analyzed the results. SZ and DK wrote the manuscript under the overall supervision by DK and all the authors approved the submission for publication.

## 7 ABBREVIATIONS

MicroRNAs: miRNAs
miR-155: microRNA-155
Pdcd4: Programmed cell death 4
*BIC*: B-cell integration gene;
AP1: Activation protein1
NFκB: nuclear factor kappa-light-chain-enhancer of activated B cells:
SAS: Tongue carcinoma cell line
FBM: Fetal buccal mucosal cell line
mRNA: Messenger RNA
qPCR: Quantitative real-time PCR
UTR: Untranslated region
WT: Wild-type
Mut: Mutant.

## REFERENCES

1. Bartel DP. 2004. MicroRNAs: genomics, biogenesis, mechanism, and function. cell 116:281–297.

2. Esquela-Kerscher A, Slack FJ. 2006. Oncomirs—microRNAs with a role in cancer. Nature Reviews Cancer 6:259.

3. Aguda BD, Kim Y, Piper-Hunter MG, Friedman A, Marsh CB. 2008. MicroRNA regulation of a cancer network: consequences of the feedback loops involving miR-17-92, E2F, and Myc. Proceedings of the National Academy of Sciences 105:19678–19683.

4. Reis PP, Tomenson M, Cervigne NK, Machado J, Jurisica I, Pintilie M, Sukhai MA, Perez-Ordonez B, Grénman R, Gilbert RW. 2010. Programmed cell death 4 loss increases tumor cell invasion and is regulated by miR-21 in oral squamous cell carcinoma. Molecular cancer 9:238.

5. O’Day E, Lal A. 2010. MicroRNAs and their target gene networks in breast cancer. Breast cancer research 12:201.

6. Chen Y, Knösel T, Kristiansen G, Pietas A, Garber ME, Matsuhashi S, Ozaki I, Petersen I. 2003. Loss of PDCD4 expression in human lung cancer correlates with tumour progression and prognosis. The Journal of Pathology: A Journal of the Pathological Society of Great Britain and Ireland 200:640–646.

7. Li J, Fu H, Xu C, Tie Y, Xing R, Zhu J, Qin Y, Sun Z, Zheng X. 2010. miR-183 inhibits TGF-β1-induced apoptosis by downregulation of PDCD4 expression in human hepatocellular carcinoma cells. BMC cancer 10:354.

8. Gao F, Zhang P, Zhou C, Li J, Wang Q, Zhu F, Ma C, Sun W, Zhang L. 2007. Frequent loss of PDCD4 expression in human glioma: possible role in the tumorigenesis of glioma. Oncology reports 17:123–128.

9. Allgayer H. 2010. Pdcd4, a colon cancer prognostic that is regulated by a microRNA. Critical reviews in oncology/hematology 73:185–191.

10. Young MR, Yang H-S, Colburn NH. 2003. Promising molecular targets for cancer prevention: AP-1, NF-κB and Pdcd4. Trends in molecular medicine 9:36–41.

11. Vikhreva P, Shepelev M, Korobko E, Korobko I. 2010. Pdcd4 tumor suppressor: properties, functions, and their application to oncology. Molekuliarnaia genetika, mikrobiologiia i virusologiia:3–11.

12. Lankat-Buttgereit B, Göke R. 2003. Programmed cell death protein 4 (pdcd4): a novel target for antineoplastic therapy? Biology of the Cell 95:515–519.

13. Yang H-S, Jansen AP, Nair R, Shibahara K, Verma AK, Cmarik JL, Colburn NH. 2001. A novel transformation suppressor, Pdcd4, inhibits AP-1 transactivation but not NF-κB or ODC transactivation. Oncogene 20.

14. Yang H-S, Jansen AP, Komar AA, Zheng X, Merrick WC, Costes S, Lockett SJ, Sonenberg N, Colburn NH. 2003. The transformation suppressor Pdcd4 is a novel eukaryotic translation initiation factor 4A binding protein that inhibits translation. Molecular and cellular biology 23:26–37.

15. Thomsen KG, Terp MG, Lund RR, Søkilde R, Elias D, Bak M, Litman T, Beck HC, Lyng MB, Ditzel HJ. 2015. miR-155, identified as anti-metastatic by global miRNA profiling of a metastasis model, inhibits cancer cell extravasation and colonization in vivo and causes significant signaling alterations. Oncotarget 6:29224.

16. Liu F, Song D, Wu Y, Liu X, Zhu J, Tang Y. 2017. MiR-155 inhibits proliferation and invasion by directly targeting PDCD 4 in non-small cell lung cancer. Thoracic cancer 8:613–619.

17. Chang SS, Jiang WW, Smith I, Poeta LM, Begum S, Glazer C, Shan S, Westra W, Sidransky D, Califano JA. 2008. MicroRNA alterations in head and neck squamous cell carcinoma. International journal of cancer 123:2791–2797.

18. Jiang S, Zhang H-W, Lu M-H, He X-H, Li Y, Gu H, Liu M-F, Wang E-D. 2010. MicroRNA-155 functions as an OncomiR in breast cancer by targeting the suppressor of cytokine signaling 1 gene. Cancer research 70:3119–3127.

19. Carinci F, Lo Muzio L, Piattelli A, Rubini C, Chiesa F, Ionna F, Palmieri A, Maiorano E, Pastore A, Laino G. 2005. Potential Markers of Tongue Tumor Progression Selected by cDNA Micro Array. International Journal of Immunopathology and Pharmacology 18:513–524.

20. Gironella M, Seux M, Xie M-J, Cano C, Tomasini R, Gommeaux J, Garcia S, Nowak J, Yeung ML, Jeang K-T. 2007. Tumor protein 53-induced nuclear protein 1 expression is repressed by miR-155, and its restoration inhibits pancreatic tumor development. Proceedings of the National Academy of Sciences 104:16170–16175.

21. Porkka KP, Pfeiffer MJ, Waltering KK, Vessella RL, Tammela TL, Visakorpi T. 2007. MicroRNA expression profiling in prostate cancer. Cancer research 67:6130–6135.

22. Calin GA, Croce CM. 2006. MicroRNA-cancer connection: the beginning of a new tale. Cancer research 66:7390–7394.

23. Faraoni I, Antonetti FR, Cardone J, Bonmassar E. 2009. miR-155 gene: a typical multifunctional microRNA. Biochimica et Biophysica Acta (BBA)-Molecular Basis of Disease 1792:497–505.

24. Zhang X, Li M, Zuo K, Li D, Ye M, Ding L, Cai H, Fu D, Fan Y, Lv Z. 2013. Upregulated miR-155 in papillary thyroid carcinoma promotes tumor growth by targeting APC and activating Wnt/β-catenin signaling. The Journal of Clinical Endocrinology & Metabolism 98:E1305–E1313.

25. Rather MI, Nagashri MN, Swamy SS, Gopinath KS, Kumar A. 2013. Oncogenic MicroRNA-155 Down-regulates Tumor Suppressor CDC73 and Promotes Oral Squamous Cell Carcinoma Cell Proliferation IMPLICATIONS FOR CANCER THERAPEUTICS. Journal of Biological Chemistry 288:608–618.

26. Marsolier J, Pineau S, Medjkane S, Perichon M, Yin Q, Flemington E, Weitzman MD, Weitzman JB. 2013. OncomiR addiction is generated by a miR-155 feedback loop in Theileria-transformed leukocytes. PLoS pathogens 9:e1003222.

27. Kong W, He L, Coppola M, Guo J, Esposito NN, Coppola D, Cheng JQ. 2010. MicroRNA-155 regulates cell survival, growth, and chemosensitivity by targeting FOXO3a in breast cancer. Journal of Biological Chemistry 285:17869–17879.

28. Louafi F, Martinez-Nunez RT, Sanchez-Elsner T. 2010. MicroRNA-155 targets SMAD2 and modulates the response of macrophages to transforming growth factor-β. Journal of Biological Chemistry 285:41328–41336.

29. Yin Q, Wang X, McBride J, Fewell C, Flemington E. 2008. B-cell receptor activation induces BIC/miR-155 expression through a conserved AP-1 element. Journal of Biological Chemistry 283:2654–2662.

30. Bohmann D, Bos TJ, Admon A, Nishimura T, Vogt PK, Tjian R. 1987. Human proto-oncogene c-jun encodes a DNA binding protein with structural and functional properties of transcription factor AP-1. Science 238:1386–1393.

31. Karin M, Liu Z-g, Zandi E. 1997. AP-1 function and regulation. Current opinion in cell biology 9:240–246.

32. Shaulian E, Karin M. 2001. AP-1 in cell proliferation and survival. Oncogene

33. Shukla GC, Singh J, Barik S. 2011. MicroRNAs: processing, maturation, target recognition and regulatory functions. Molecular and cellular pharmacology 3:83.

34. Franken NA, Rodermond HM, Stap J, Haveman J, Van Bree C. 2006. Clonogenic assay of cells in vitro. Nature protocols 1:2315.

35. Borowicz S, Van Scoyk M, Avasarala S, Mk KR, Tauler J, Bikkavilli RK, Winn RA. 2014. The soft agar colony formation assay. Journal of visualized experiments: JoVE:e51998–e51998.

36. Göke R, Barth P, Schmidt A, Samans B, Lankat-Buttgereit B. 2004. Programmed cell death protein 4 suppresses CDK1/cdc2 via induction of p21 Waf1/Cip1. American Journal of Physiology-Cell Physiology 287:C1541–C1546.

37. Sun Q, Miao J, Luo J, Yuan Q, Cao H, Su W, Zhou Y, Jiang L, Fang L, Dai C. 2018. The feedback loop between miR-21, PDCD4 and AP-1 functions as a driving force for renal fibrogenesis. J Cell Sci 131:jcs202317.

38. Zhang Y, Shen W-L, Shi M-L, Zhang L-Z, Zhang Z, Li P, Xing L-Y, Luo F-Y, Sun Q, Zheng X-F. 2015. Involvement of aberrant miR-139/Jun feedback loop in human gastric cancer. Biochimica et Biophysica Acta (BBA)-Molecular Cell Research 1853:481–488.

39. Moles R. 2017. MicroRNAs-based Therapy: A Novel and Promising Strategy for Cancer Treatment. MicroRNA (Shariqah, United Arab Emirates) 6:102.

40. Zhu Q, Wang Z, Hu Y, Li J, Li X, Zhou L, Huang Y. 2012. miR-21 promotes migration and invasion by the miR-21-PDCD4-AP-1 feedback loop in human hepatocellular carcinoma. Oncology reports 27:1660–1668.

41. Frankel LB, Christoffersen NR, Jacobsen A, Lindow M, Krogh A, Lund AH. 2008. Programmed cell death 4 (PDCD4) is an important functional target of the microRNA miR-21 in breast cancer cells. Journal of Biological Chemistry 283:1026–1033.

42. Asangani I, Rasheed S, Nikolova D, Leupold J, Colburn N, Post S, Allgayer H. 2008. MicroRNA-21 (miR-21) post-transcriptionally downregulates tumor suppressor Pdcd4 and stimulates invasion, intravasation and metastasis in colorectal cancer. Oncogene 27:2128–2136.

43. Gu W, Gao T, Shen J, Sun Y, Zheng X, Wang J, Ma J, Hu X-Y, Li J, Hu M-J. 2014. MicroRNA-183 inhibits apoptosis and promotes proliferation and invasion of gastric cancer cells by targeting PDCD4. International journal of clinical and experimental medicine 7:2519.

44. Ma Q-q, Huang J-t, Xiong Y-g, Yang X-y, Han R, Zhu W-w. 2017. MicroRNA-96 Regulates Apoptosis by Targeting PDCD4 in Human Glioma Cells. Technology in cancer research & treatment 16:92–98.

45. Zhang X, Gee H, Rose B, Lee CS, Clark J, Elliott M, Gamble JR, Cairns MJ, Harris A, Khoury S. 2016. Regulation of the tumour suppressor PDCD4 by miR-499 and miR-21 in oropharyngeal cancers. BMC cancer 16:86.

46. Ameyar M, Wisniewska M, Weitzman J. 2003. A role for AP-1 in apoptosis: the case for and against. Biochimie 85:747–752.

